# Genome-wide association analysis of lifetime cannabis use (N=184,765) identifies new risk loci, genetic overlap with mental health, and a causal influence of schizophrenia on cannabis use

**DOI:** 10.1101/234294

**Authors:** Joëlle A. Pasman, Karin J.H. Verweij, Zachary Gerring, Sven Stringer, Sandra Sanchez-Roige, Jorien L. Treur, Abdel Abdellaoui, Michel G. Nivard, Bart M.L. Baselmans, Jue-Sheng Ong, Hill F. Ip, Matthijs D. van der Zee, Meike Bartels, Felix R. Day, Pierre Fontanillas, Sarah L. Elson, the 23andMe Research Team, Harriet de Wit, Lea K. Davis, James MacKillop, International Cannabis Consortium, Jaime L. Derringer, Susan J.T. Branje, Catharina A. Hartman, Andrew C. Heath, Pol A.C. van Lier, Pamela A.F. Madden, Reedik Mägi, Wim Meeus, Grant W. Montgomery, A.J. Oldehinkel, Zdenka Pausova, Josep A. Ramos-Quiroga, Tomas Paus, Marta Ribases, Jaakko Kaprio, Marco P.M. Boks, Jordana T. Bell, Tim D. Spector, Joel Gelernter, Dorret I. Boomsma, Nicholas G. Martin, Stuart MacGregor, John R.B. Perry, Abraham A. Palmer, Danielle Posthuma, Marcus R. Munafò, Nathan A. Gillespie, Eske M. Derks, Jacqueline M. Vink

**Affiliations:** Behavioural Science Institute, Radboud University, Nijmegen, The Netherlands; Genetic Epidemiology, Statistical Genetics, and Translational Neurogenomics laboratories, QIMR Berghofer Medical Research Institute, Brisbane, Queensland, Australia; Department of Complex Trait Genetics, Center for Neurogenomics and Cognitive Research, Vrije Universiteit Amsterdam, Amsterdam, The Netherlands; University of California San Diego, Department of Psychiatry, La Jolla, United States of America; MRC Integrative Epidemiology Unit (IEU), University of Bristol, Bristol, United Kingdom; Department of Biological Psychology/Netherlands Twin Register, Vrije Universiteit Amsterdam, Amsterdam, The Netherlands; MRC Epidemiology Unit, University of Cambridge School of Clinical Medicine, Institute of Metabolic Science, Cambridge Biomedical Campus, Cambridge, United Kingdom; 23andMe, Inc., Mountain View, United States of America; Department of Psychiatry and Behavioral Neuroscience, University of Chicago, Chicago, United States of America; Vanderbilt Genetics Institute; Division of Genetic Medicine, Department of Medicine, Vanderbilt University, Nashville, United States of America; Peter Boris Centre for Addictions Research and Michael G. DeGroote Centre for Medicinal Cannabis Research, McMaster University/St. Joseph's Healthcare Hamilton, Hamilton, Canada; Department of Psychology, University of Illinois Urbana-Champaign, Champaign, United States of America; Department of Youth and Family, Utrecht University, Utrecht, The Netherlands; Department of Psychiatry, Interdisciplinary Center Psychopathology and Emotion Regulation, University of Groningen, University Medical Center Groningen, Groningen, The Netherlands; Department of Psychiatry, Washington University School of Medicine, St. Louis (MO), United States of America; Department of Developmental Psychology and EMGO Institute for Health and Care Research, Vrije Universiteit Amsterdam, Amsterdam, The Netherlands; Estonian Genome Center, University of Tartu, Tartu, Estonia; Institute for Molecular Bioscience, The University of Queensland, Brisbane, Queensland, Australia; Hospital for Sick Children, Toronto, Canada; Psychiatric Genetics Unit, Group of Psychiatry, Mental Health and Addiction, Vall d’Hebron Research Institute (VHIR), Universitat Autònoma de Barcelona, Spain; Department of Psychiatry, Hospital Universitari Vall d’Hebron, Barcelona, Spain; Biomedical Network Research Centre on Mental Health (CIBERSAM), Instituto de Salud Carlos III, Barcelona, Spain; Department of Psychiatry and Legal Medicine, Universitat Autònoma de Barcelona, Barcelona, Spain; Rotman Research Institute, Baycrest, Toronto, Canada; Departments of Psychology and Psychiatry, University of Toronto, Toronto, Canada; Institute for Molecular Medicine Finland FIMM, HiLIFE Unit, University of Helsinki, Helsinki, Finland; Brain Center Rudolf Magnus, Department of Psychiatry, University Medical Center Utrecht, Utrecht, The Netherlands; Department of Twin Research and Genetic Epidemiology, King’s College London, London, United Kingdom; Department of Psychiatry, Yale University School of Medicine, New Haven (CT), United States of America; University of California San Diego, Institute for Genomic Medicine, La Jolla, United States of America; UK Centre for Tobacco and Alcohol Studies and School of Experimental Psychology, University of Bristol, Bristol, United Kingdom; Department of Psychiatry, Virginia Institute for Psychiatric and Behavior Genetics, Virginia Commonwealth University, Richmond, Virginia, United States of America

## Abstract

Cannabis use is a heritable trait [1] that has been associated with adverse mental health outcomes. To identify risk variants and improve our knowledge of the genetic etiology of cannabis use, we performed the largest genome-wide association study (GWAS) meta-analysis for lifetime cannabis use (N=184,765) to date. We identified 4 independent loci containing genome-wide significant SNP associations. Gene-based tests revealed 29 genome-wide significant genes located in these 4 loci and 8 additional regions. All SNPs combined explained 10% of the variance in lifetime cannabis use. The most significantly associated gene, *CADM2*, has previously been associated with substance use and risk-taking phenotypes [2–4]. We used S-PrediXcan to explore gene expression levels and found 11 unique eGenes. LD-score regression uncovered genetic correlations with smoking, alcohol use and mental health outcomes, including schizophrenia and bipolar disorder. Mendelian randomisation analysis provided evidence for a causal positive influence of schizophrenia risk on lifetime cannabis use.

---

We performed the largest GWAS of lifetime cannabis use (having ever tried cannabis) to date. We meta-analysed 3 GWASs (International Cannabis Consortium [ICC,] N=35,297; 23andMe, N=22,683; UK-Biobank, N=126,785) with a combined sample size of 184,765 individuals, a five-fold increase compared to the previous largest GWAS for lifetime cannabis use [5]. The meta-analysis resulted in 646 genome-wide significant SNP associations located in 4 independent (linkage disequilibrium [LD] R^2^<0.1, window size 250 kb) regions on chromosomes 3, 8, 11, and 16 (Table 1, Figure 1, and Supplementary Table S1). The most strongly associated marker was an intronic variant of *CADM2* on chromosome 3 (rs2875907, *p*=2.66e-15). Other hits were located in *ZNF704, NCAM1*, and *RABEP2/ATP2A1* (Figure 2). All tested SNPs combined explained 10% (h^2^_SNP_=0.10, SE=0.01) of the individual differences in lifetime cannabis use; approximately 25% of twin-based heritability estimates [1]. Supplementary Figure S1-3 and Table S2 provide information on results of the individual GWASs.

**Table 1.**
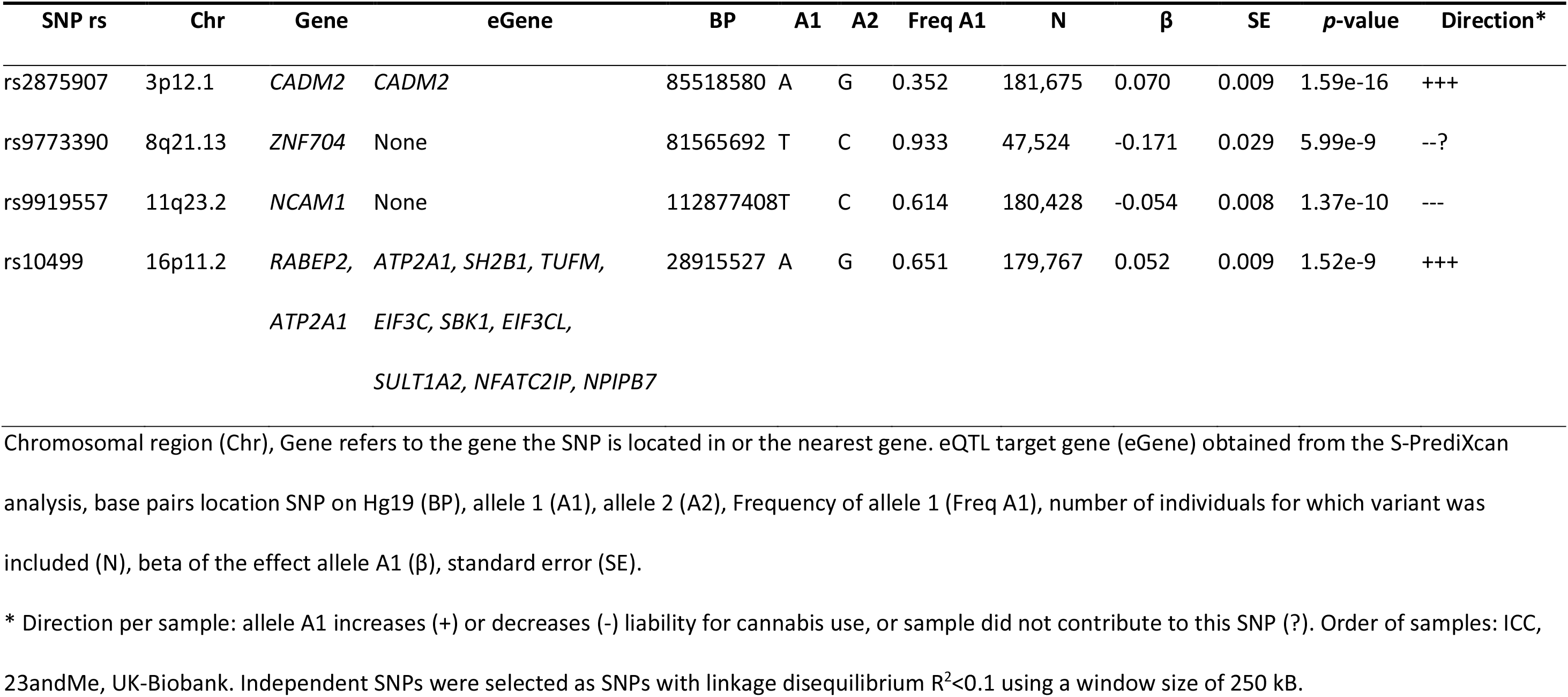
Association results of 4 independent SNPs that are significantly associated with lifetime cannabis use.

**Figure 1.**
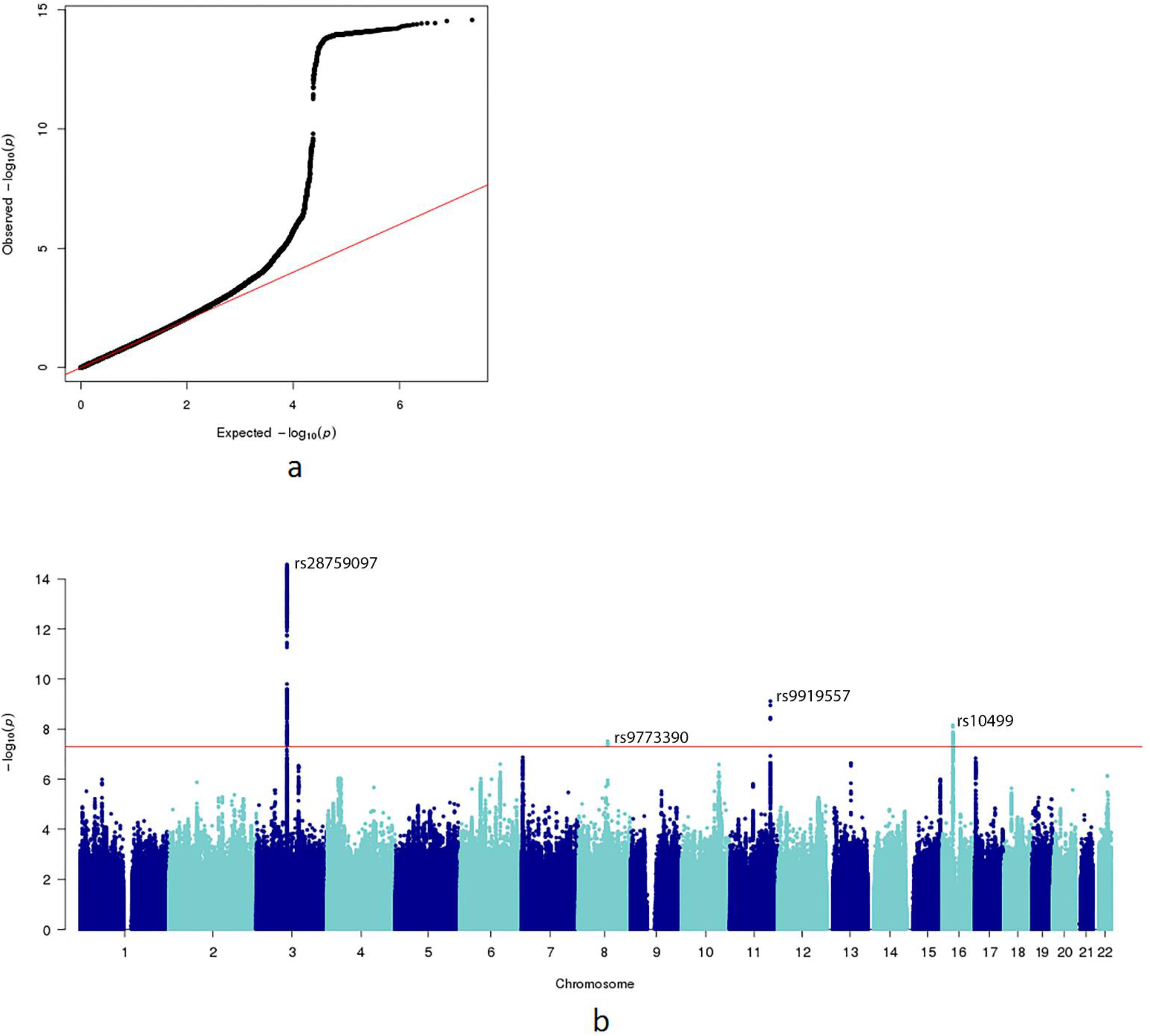
A) QQ-plot of the distribution of the −log_10_(p-values) observed for the SNP associations with lifetime cannabis use against those expected under the null hypothesis. There was no evidence for stratification effects (LD score regression b_0_=0.90, SE=0.007). B) Manhattan plot for the SNP-based GWAS meta-analysis. The SNP with the smallest p-value per genome-wide significant locus is annotated. The red line represents the genome-wide significance threshold of *p*<5e-08.

**Figure 2.**
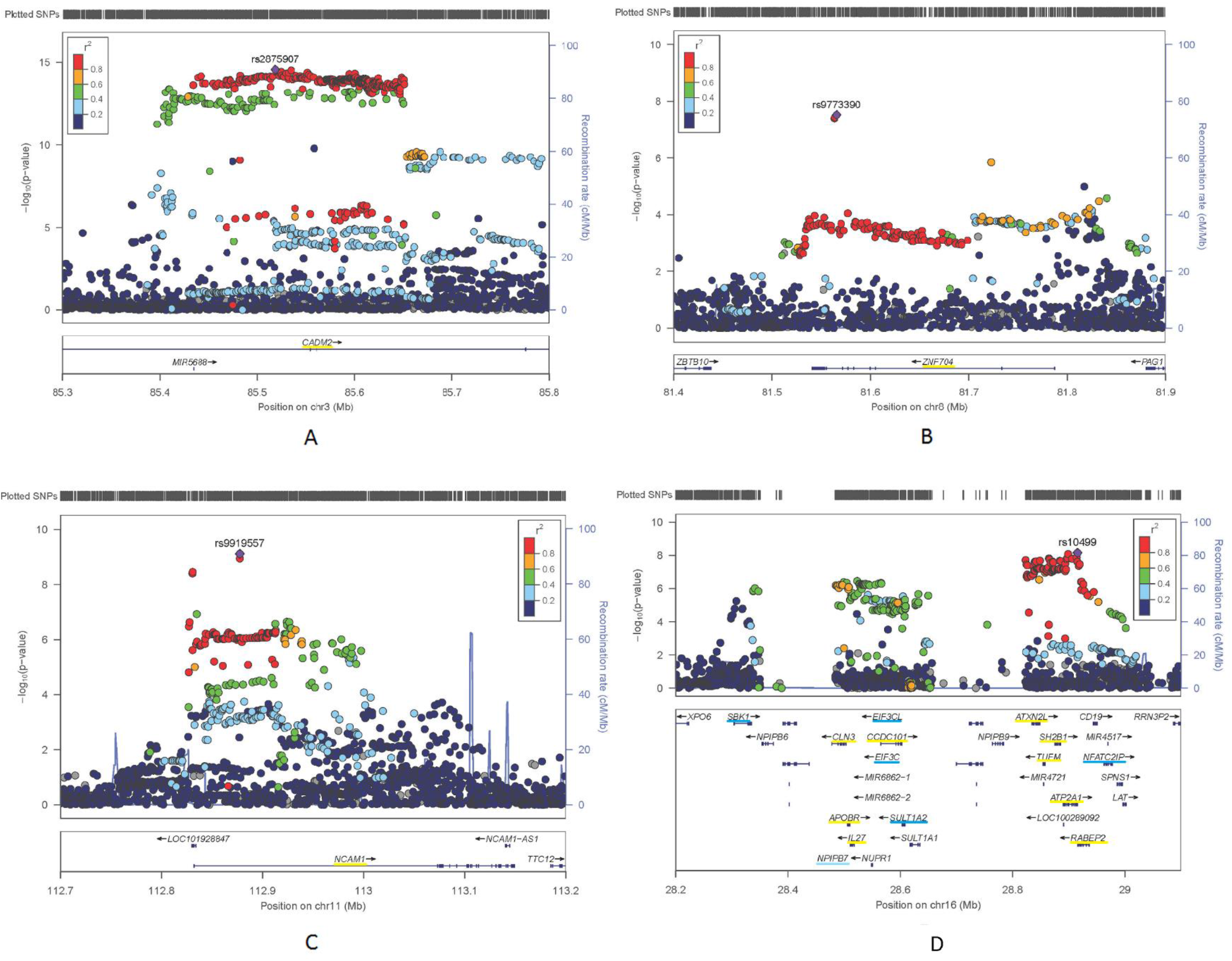
Regional plots for the genome-wide significant hits, with A) lead SNP rs2875907 on chromosome 3, B) rs9773390 on chromosome 8, C) rs9919557 on chromosome 11, and D) rs10499 on chromosome 16. Underlined in yellow genes that were identified in the gene-based test; in blue genes that were identified in the S-PrediXcan analysis only (in light blue pseudo genes).

Gene-based tests of associations in MAGMA [6] identified 29 genes significantly associated with lifetime cannabis use (Figure 3, Table 2, and Supplementary Figure S4-5). These were located in the 4 regions identified in the SNP-based analysis and in 8 putatively novel regions. *CADM2* and *NCAM1*, both previously identified in the original ICC meta-analysis [5], were among the strongest hits. The *CADM2* gene (Cell Adhesion Molecule 2) is a synaptic cell adhesion molecule and is part of the immunoglobulin superfamily. Interestingly, *CADM2* has previously been identified in GWAS of other behavioural phenotypes, including alcohol consumption [2], processing speed [7], and number of offspring and risk-taking behavior [4]. A large-scale phenome-wide scan showed that *CADM2* was associated with various personality traits, with the risk variant being associated with e.g. reduced anxiety, neuroticism and conscientiousness, and increased risk-taking [3]. Taken together, these findings suggest that risk variants in *CADM2* are associated with a broad profile of a risk-taking, optimistic, and care-free personality [3]. Cannabis use has previously been associated with these personality traits, including high levels of impulsivity and novelty seeking [8, 9].

**Table 2.** Genes identified in the MAGMA and/ or S-PrediXcan analyses. Gene-based test parameters are provided for genes with a *p*-value below 2.74e-06 in the MAGMA test (*p*<0.05 with Bonferroni correction for the 18,269 tested genes). If genes were significant in the S-PrediXcan analysis only the corresponding S-PrediXcan parameters were included and the row was hightlighted (gray).

| Locus | Top genes | BP start | BP stop | #SNPs | Z | $p$-value |
| --- | --- | --- | --- | --- | --- | --- |
| 2p12 | *LRRTM4* | 76969849 | 77754502 | 3561 | 4.72 | 1.15e-06 |
| 3p12.1 | *CADM2* | 85003133 | 86128579 | 4262 | 8.47 | 1.28e-17 |
| 6p12.1 | *BEND6* | 56814773 | 56897450 | 220 | 5.86 | 1.83e-07 |
|  | *KIAA1586* | 56906343 | 56925023 | 30 | 4.80 | 7.89e-07 |
|  | *BAG2* | 57032104 | 57055013 | 36 | 4.64 | 1.71e-06 |
|  | *RAB23* | 57046790 | 57092112 | 73 | 5.60 | 1.06e-08 |
| 6q25.3 | *ARID1B* | 157093980 | 157536913 | 1324 | 4.75 | 1.01e-06 |
| 8q21.13 | *ZNF704* | 81535686 | 81792016 | 526 | 4.75 | 1.01e-06 |
| 10q24.32 | *AS3MT* | 104624183 | 104666656 | 151 | 5.29 | 6.23e-08 |
|  | *CNNM2* | 104673075 | 104843344 | 516 | 5.04 | 2.31e-07 |
|  | *NT5C2* | 104842774 | 104958063 | 347 | 4.62 | 1.92e-06 |
| 11q23.2 | *NCAM1* | 112826969 | 113154158 | 1234 | 6.19 | 2.94e-10 |
| 12q24.12 | *ACAD10* | 112118857 | 112199911 | 130 | 4.90 | 4.91e-07 |
|  | *ALDH2* | 112199691 | 112252789 | 88 | 4.78 | 8.97e-07 |
|  | *MAPKAPK5* | 112275032 | 112336228 | 168 | 4.63 | 1.87e-06 |
|  | *TMEM116* | 112364086 | 112456023 | 207 | 4.66 | 1.61e-06 |
| 16q12.1 | *SBK1* | 28303840 | 28335170 | 16 | 5.57 | 2.51e-08 |
|  | *NPIPB7* | 28467693 | 28481868 | 15 | 5.28 | 1.30e-07 |
|  | *CLN3* | 28483600 | 28510897 | 53 | 5.41 | 3.19e-08 |
|  | *APOBR* | 28500970 | 28515291 | 37 | 5.37 | 4.02e-08 |
|  | *IL27* | 28505683 | 28523155 | 37 | 5.40 | 3.42e-08 |
|  | *CCDC101* | 28560249 | 28608111 | 143 | 4.68 | 1.41e-06 |
|  | *SULT1A2* | 28603264 | 28608391 | 9 | 5.38 | 7.56e-08 |
|  | *EIF3C* | 28722782 | 28747053 | 6 | 5.65 | 1.59e-08 |
|  | *EIF3CL* | 28722785 | 28747053 | 16 | 5.51 | 3.66e-08 |
|  | *ATXN2L* | 28829369 | 28853558 | 63 | 5.63 | 8.79e-09 |
|  | *TUFM* | 28848732 | 28862729 | 34 | 5.60 | 1.05e-08 |
|  | *SH2B1* | 28867939 | 28890534 | 52 | 5.50 | 1.91e-08 |
|  | *ATP2A1* | 28884192 | 28920830 | 65 | 5.63 | 9.28e-09 |
|  | *NFATC2IP* | 28962318 | 28977767 | 14 | 5.32 | 1.03e-07 |
|  | *RABEP2* | 28910742 | 28942339 | 58 | 5.13 | 1.54e-07 |
| 17p13.3 | *SRR* | 2202244 | 2233553 | 85 | 5.39 | 3.44e-08 |
|  | *TSR1* | 2220972 | 2245678 | 68 | 5.20 | 9.84e-08 |
| 18q11.2 | *C18orf8* | 21078434 | 21118311 | 98 | 4.98 | 3.13e-07 |
|  | *NPC1* | 21081148 | 21171581 | 220 | 5.01 | 2.78e-07 |

Gene number corresponding to regional plots (ID), gene name (Symbol), Chromosome (Chr), location in base pairs (hg19) at beginning and end of gene (BP start/stop), number of SNPs included in the gene (#SNPs), test-statistic for the test of association (Z).

**Figure 3.**
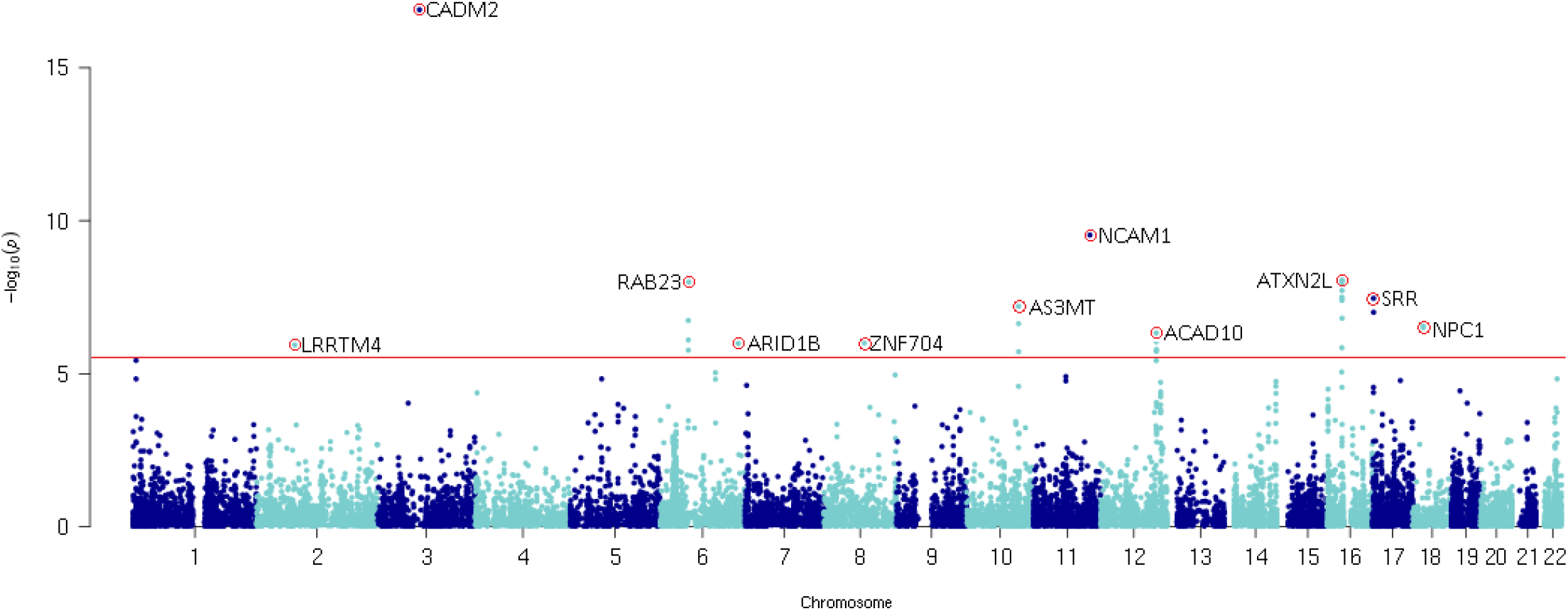
Manhattan plot for the gene-based test of association. The red line represents the genome-wide significance threshold of *p*<2.74e-06.

*NCAM1* (Neural Cell Adhesion Molecule 1) also encodes a cell adhesion protein and is member of the immunoglobulin superfamily. The encoded protein is involved in cell-matrix interactions and cell differentiation during development [10]. *NCAM1* is located in the *NCAM1-TTC12-ANKK1-DRD2* gene cluster, which is related to neurogenesis and dopaminergic neurotransmission. This gene cluster has been associated with smoking, alcohol use, and illicit drug use [11–14] and has been implicated in psychiatric disorders, such as schizophrenia and mood disorders [15, 16].

Putatively novel findings in both the SNP- and gene-based test were the ZNF704 region at chromosome 8, about which little is known, and *RABEP2/ATP2A1*, located in an interesting region on chromosome 16 (see below). Several of the 29 top genes have previously shown an association with schizophrenia (e.g., *TUFM, NCAM1*), BMI or obesity, alcohol use (e.g. *ALDH2*), intelligence and cognitive performance, and externalizing and impulsive phenotypes (Supplementary Table S3). At the phenotypic level, associations between cannabis use and psychiatric disorders [17], use of other substances [8], personality [18], and educational attainment [19] are well-established.

S-PrediXcan analysis, aimed at identifying genes with differential expression levels in cannabis users versus non-users [eGenes, 20], largely confirmed SNP- and gene-based findings. S-PrediXcan revealed 51 Bonferroni-corrected significant associations across tissues (Supplementary Tables S3 and S4) targeting 11 unique eGenes. Five eGenes were also significant in the gene-based tests, whereas 6 were novel. For eGenes identified in multiple tissues, directions of effects were consistent across tissues (Supplementary Table S4). Again, the top finding was *CADM2*; genetic variants associated with increased liability to use cannabis are predicted to upregulate expression levels of *CADM2* in 5 (non-brain) tissues, including whole blood. Of note, although *CADM2* is expressed more widely in brain compared with other tissues (Supplementary Figure S6), rs2875907 regulates the expression of *CADM2* only in non-brain tissues (Supplementary Figure S7). Exploration of S-PrediXcan results in UK-Biobank data (https://imlab.shinyapps.io/gene2pheno_ukb_neale/) showed that *CADM2* expression is significantly associated with multiple traits, including increased risk-taking and BMI, and reduced feelings of anxiety.

As did the SNP- and gene-based tests, the S-PrediXcan analysis detected a strong signal in a single high-LD region at 16p11.2. Deletions and duplications in this region were previously reported to be associated with autism and schizophrenia [21, 22], while a common 16p11.2 inversion underlies susceptibility to asthma and obesity [23]. The inversion explains a substantial proportion of variability in expression of the eGenes, including *TUFM* and *SH2B1* [23]. Due to high LD in this region and high levels of co-expression of the eGenes, follow-up studies will be needed to determine which gene(s) are functionally driving the association with cannabis use.

Using our GWAS results and those of other studies, we estimated the genetic correlation of lifetime cannabis use with 25 other traits of interest with LD score regression. Fourteen traits were significantly genetically correlated with lifetime cannabis use, after correction for multiple testing (Figure 4 and Supplementary Table S5). Positive genetic correlations were found with substance use phenotypes, including smoking and alcohol use and dependence, as well as with mental health phenotypes, including ADHD and schizophrenia. Furthermore, positive genetic correlations were found with risk-taking behaviour, openness to experience, and educational attainment, as well as a negative correlation with conscientiousness. The broad range of correlations suggests that genetic liability to cannabis use should be viewed in a larger context of personality and mental health traits. Specifically, the substantial genetic correlations with risk-taking behaviour and openness to experience may indicate liability to start using cannabis is an indication of one’s personality.

**Figure 4.**
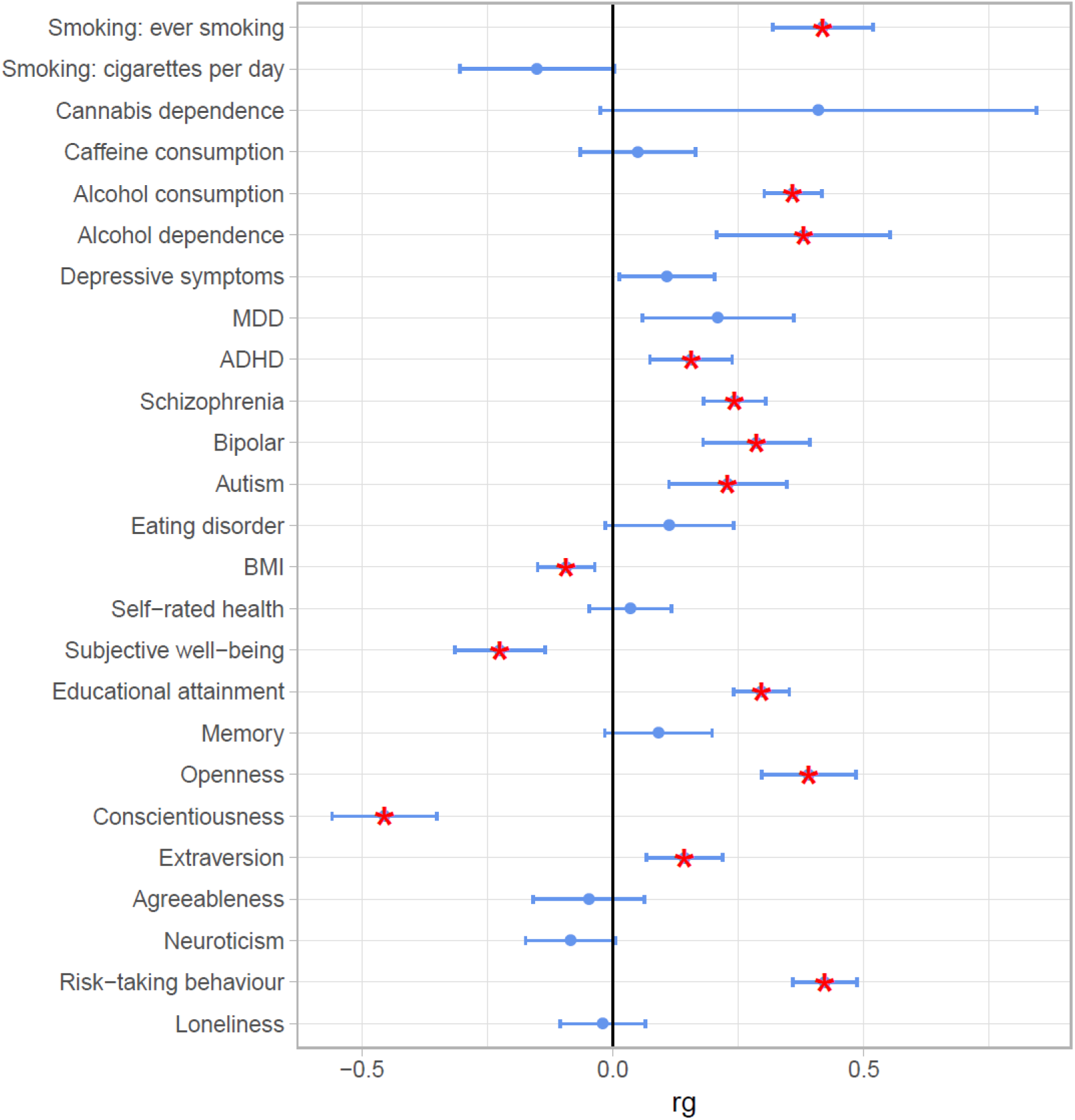
Genetic overlap between lifetime cannabis use and other phenotypes. Error bars represent 95% confidence intervals and asterisks indicate significant associations after correction for multiple testing.

The relationship between cannabis use and schizophrenia has been subject of intensive research and debate. It has long been established that cannabis use is higher in schizophrenia patients [24, 25]. A substantial body of evidence supports the hypothesis that cannabis use increases the risk for developing schizophrenia [26], but other hypotheses (i.e. schizophrenia increases use of cannabis, or the association is due to (genetic) pleiotropy) have also been posed. Our results confirm previous findings [27, 28] that genetic risk factors for cannabis use and schizophrenia are positively correlated (r_g_=0.24, SE=0.03, p<0.01). However, since a genetic correlation does not provide insight in the direction of causation, we performed bi-directional two-sample Mendelian randomisation (MR) analysis [29] to examine whether there is evidence for a causal relationship from cannabis use to schizophrenia and vice versa.

We found no clear evidence for a causal influence of lifetime cannabis use on schizophrenia risk; the Inverse Variance Weighted [IVW] regression odds ratios were 1.01 (95% CIs 0.70-1.45, *p*=0.97) and 1.03 (95% CIs 0.89-1.19, *p*=0.70) for the 5e-08 and 1e-05 p-value thresholds for SNP inclusions, respectively. We did find evidence for a causal positive influence of schizophrenia risk on lifetime cannabis use (IVW regression OR=1.16, 95% CIs 1.06-1.27, *p*<0.01) (Table 3; Supplementary Tables S6-S9 and Supplementary Figures S8-S9 for details). We performed 4 sensitivity analyses to determine the robustness of this finding; these analyses provided a consistent pattern of effect sizes (with the exception of the MR-Egger analysis) supporting the causal effect from schizophrenia to cannabis use, albeit with weaker statistical evidence (Table 3). Moreover, the MR-Egger intercept was not significant (Supplementary Table S9), indicating no evidence for pleiotropy [30]. Note that while these methods allow us to infer causality, they do not provide interpretable estimates of the magnitude of the causal effect, as the phenotypes were measured on a logistic scale.

**Table 3.** Bidirectional two-sample Mendelian randomisation analyses between lifetime cannabis use and schizophrenia.

|  | *Cannabis-Schizophrenia* |  |  |  | *Cannabis-Schizophrenia* |  |  |  | *Schizophrenia-Cannabis* |  |  |  |
| --- | --- | --- | --- | --- | --- | --- | --- | --- | --- | --- | --- | --- |
|  | *($p < 5e-08$, 4 SNPs)* |  |  |  | *($p < 1e-05$, 37 SNPs)* |  |  |  | *($p < 5e-08$, 110 SNPs)* |  |  |  |
|  | **B** | **SE (B)** | **OR** | ***p*** | **B** | **SE (B)** | **OR** | ***p*** | **B** | **SE (B)** | **OR** | ***p*** |
| **IVW** | 0.007 | 0.187 | 1.01 | 0.971 | 0.029 | 0.075 | 1.03 | 0.695 | 0.150 | 0.046 | 1.16 | 0.001 |
| **Weighted median** | -0.085 | 0.100 | 0.92 | 0.398 | -0.004 | 0.061 | 1.00 | 0.942 | 0.164 | 0.051 | 1.18 | 0.001 |
| **MR-Egger** | -0.098 | 0.544 | 0.91 | 0.873 | 0.013 | 0.183 | 1.01 | 0.943 | 0.059 | 0.212 | 1.06 | 0.817 |
| **Simple Mode** | -0.208 | 0.123 | 0.81 | 0.190 | -0.109 | 0.119 | 0.90 | 0.366 | 0.321 | 0.173 | 1.38 | 0.066 |
| **Weighted Mode** | -0.101 | 0.107 | 0.90 | 0.414 | -0.050 | 0.084 | 0.95 | 0.559 | 0.315 | 0.185 | 1.37 | 0.091 |

Inverse Variance Weighted regression analysis (IVW); risk coefficient representing the change in outcome for a one-unit increase in the exposure variable
(B); standard error of the B (SE (B)); odds ratios represent the odds of schizophrenia for lifetime cannabis users versus non-users (when cannabis is the exposure) or the odds of lifetime cannabis use for those with a schizophrenia diagnosis versus those without (when schizophrenia is the exposure) (OR); *p*-value (*p*).

Two previous two-sample MR studies investigated the link between lifetime cannabis use and schizophrenia. The first only tested causal effects from cannabis use to schizophrenia and found evidence for causality [31], in contrast to our findings. The second study tested bi-directional effects with genetic instruments more similar to ours and found weak evidence for a causal effect of cannabis use to schizophrenia and much stronger evidence for a causal effect in the other direction [32]. Our results reinforce this latter finding, suggesting that genetic risk for schizophrenia causally contributes to an increased liability to use cannabis. A possibility is that individuals at risk for developing schizophrenia experience prodromal symptoms or negative affect that make them more likely to start using cannabis to cope or self-medicate [33]. The lack of evidence of a causal influence of cannabis use on schizophrenia may be due to the lower power of the instrumental variables. The instrumental variable based on schizophrenia SNPs explained 3.38% of variance in liability to schizophrenia. For cannabis use, the genetic instruments explained 0.63% and 0.12% of the variance in cannabis use for SNPs included with *p*<1e-05 and *p*<5e-08, respectively.

Our GWAS of lifetime cannabis use, which is the largest to date, revealed significant associations in 12 putatively novel regions. Among these, the most promising candidates for future functional studies are *CADM2, NCAM1*, the *ZNF704* region, and multiple genes located at 16p11.2. Our findings further indicated a causal influence of schizophrenia on cannabis use and substantial genetic overlap between cannabis use and use of other substances, mental health, and personality traits, such as risk-taking and extraversion.

## Online Methods

### Samples

Data from 3 sources were obtained: ICC, 23andMe and UK-Biobank (total N=184,765). We used existing GWAS summary statistics from the **ICC**, based on data from 35,297 individuals of European ancestry from 16 cohorts from Northern America, Europe, and Australia [5]. The overall sample included 55.5% females and the age ranged between 16 and 87 years with a mean of 35.7 years. An average of 42.8% of the individuals had used cannabis during their lifetime. The second set of results is derived from the personal genetics company **23andMe** Inc. Data were available for 22,683 individuals of European Ancestry who provided informed consent and answered surveys online according to a human subjects protocol approved by Ethical & Independent Review Services, a private institutional review board. The sample included 55.3% females and the age ranged between 18 and 94 years with a mean of 54.0 years. Within the sample, 43.2% had used cannabis during their lifetime. The third sample is obtained from **UK-Biobank**. Data were available for 126,785 individuals of European ancestry. The sample included 56.3% females and the age ranged between 39 and 72 years with a mean of 55.0 years. Within the sample, 22.3% had used cannabis during their lifetime.

### Phenotype and covariates

For all participants, self-report data were available on whether the participant had ever used cannabis during their lifetime: yes (1) versus no (0). Measurement instruments and phrasing of the questions about lifetime cannabis use differed across the samples. For the ICC study this has been described for each cohort in the original paper [5]. As part of their online questionnaire, 23andMe used the following phrase to examine lifetime cannabis use: ‘Have you ever in your life used the following: Marijuana?’. The UK-Biobank – as part of an online follow-up questionnaire - asked: ‘Have you taken CANNABIS (marijuana, grass, hash, ganja, blow, draw, skunk, weed, spliff, dope), even if it was a long time ago?’.

### Genotyping and imputation

Genotyping was performed on various genotyping platforms and standard quality control checks were performed prior to imputation. Genotype data were imputed using the 1000 Genomes phase 1 release reference set [34] for ICC and 23andMe, and the Haplotype Reference Consortium reference set [35] for the UK-Biobank sample. Information about samples, genotyping, and imputation is summarized in Supplementary Table S10. After quality control, the ICC sample comprised 35,297 individuals and 6,643,927SNPs, 23andMe 22,683 individuals and 7,837,888 SNPs, and the UK-Biobank sample 126,785 individuals and 10,827,718 SNPs.

### Genome-wide association analyses

We conducted the GWAS in 23andMe and UK-Biobank samples separately. Associations between the binary phenotype and SNPs were tested using a logistic regression model accounting for the effects of sex, age, ancestry, and genotype batch (and age^2^ in the UK-Biobank sample). The GWAS for UK-Biobank were performed in PLINK 1.9 [36] and for 23andMe in an internally developed pipeline. We then meta-analysed the GWAS results from ICC, 23andMe, and UK-Biobank. Prior to conducting the meta-analysis, additional quality control of the summary statistics of each study was conducted in EasyQC [37]. Because of varying GWAS methods and sample characteristics (see Supplementary Table S10), slightly different quality control criteria were used for the 3 samples (see Supplementary Table S11). All 3 samples were aligned with the Haplotype Reference Consortium panel using the EasyQC R-package [37], in order to ensure that RS-numbers and chromosome-basepair positions referred to the exact same variants and to correct for strand effects. Variants were deleted if they had a minor allele frequency diverging more than 0.15 from that in the reference panel.

We applied genomic control to the 3 GWAS files prior to meta-analysis to ensure none of the samples contributed disproportionately to the meta-analysis results [38]. Inflation due to stratification was estimated using LD-score regression; the intercept was used to correct the test statistics (b_0_= 1.003, SE=0.007 for ICC, b_0_=1.004, SE=0.006) for 23andMe, b_0_=1.022, SE=0.007 for UK-Biobank). We then performed a fixed effects meta-analysis based on effect sizes (log odds ratios (OR)) and standard errors in METAL [39]. We applied the conventional p-value threshold of 5e-08 as an indication of genome-wide significance. The meta-analysis was performed on 11,696,151 SNPs that passed quality control. The combined sample size of the meta-analysis was 184,764 individuals, but note that the sample size varies per SNP due to differential missingness across samples.

Regional plots were created using LocusZoom [40], with varying window size for optimal visualization (see Supplementary Figure S5).

### Gene-based test of association

Testing associations on the level of protein-coding genes is more biologically meaningful and more powerful (lower multiple testing burden) than testing solely on the level of SNPs. Gene-based analysis was used to test associations for aggregates of variants in protein-coding genes. The analysis was conducted in MAGMA (v 1.6) [6], which uses the 1000 Genomes reference-panel (phase 3, 2012) to control for LD. SNPs were mapped to genes if they were located in or within 5 kb from the gene, such that 4,760,663 SNPs (41%) could be mapped to at least one of 18,269 protein-coding genes in the reference panel. The significance threshold was set at a Bonferroni corrected p<0.05 (0.05/18,269=2.74e-06).

### SNP-based heritability analysis

The proportion of variance in liability to cannabis use that could be explained by the aggregated effect of all SNPs (h^2^_SNPs_) was estimated using LD-Score Regression analysis [41]. The method is based on the premise that an estimated SNP effect-size includes effects of all SNPs in linkage disequilibrium (LD) with that SNP. A SNP that tags many other SNPs will have a higher probability of tagging a causal genetic variant compared to a SNP that tags few other SNPs. The LD score measures the amount of genetic variation tagged by a SNP within a specific population. Accordingly, assuming a trait with a polygenic architecture, SNPs with a higher LD-score have on average stronger effect sizes than SNPs with lower LD-scores. When regressing the effect size from the association analysis against the LD score for each SNP, the slope of the regression line provides an estimate of the proportion of variance accounted for by all analysed SNPs [41]. For this analysis, we included 1,179,898 SNPs that were present in all cohorts and the HapMap 3 reference panel. Standard LD scores were used as provided by Bulik-Sullivan et al. [41] based on the Hapmap 3 reference panel, restricted to European populations.

### Identification of genes with differential expression levels between cannabis users and non-users

We used summary S-PrediXcan to integrate eQTL information with summary statistics from the lifetime cannabis use GWAS meta-analysis to identify eGenes (i.e., genes of which genetically predicted expression levels are associated with cannabis use [20]). Briefly, S-PrediXcan estimates gene expression weights by training a linear prediction model in samples with both gene expression and SNP genotype data. The weights are then used to predict gene expression from GWAS summary statistics, while incorporating the variance and co-variance of SNPs from an LD reference panel. We used expression weights for 44 tissues from the GTEx Project (V6p) and the DGN whole blood cohort, generated by Gamazon et al. [42] and LD information from the 1000 Genome Project Phase 3 [43]. These data were processed with beta values and standard errors from the lifetime cannabis use GWAS meta-analysis to estimate the expression-GWAS association statistic. We used a transcriptome-wide significant threshold of *p*<2.53e-07, which is the Bonferroni corrected threshold when adjusting for all tissues and genes (i.e., 197,680 gene-based tests in the GTEx and DGN reference sets).

We used the GTEXPortal [https://www.gtexportal.org/home/;GTExAnalysisReleaseV7;44] to obtain gene expression levels of *CADM2* across tissues. We used the same portal to plot a multi-tissue eQTL comparison of the top SNP rs2875907. The multi-tissue eQTL plot shows both the single-tissue eQTL p-value and the multi-tissue posterior probability from METASOFT [45].

### Genetic correlation with use of other substances and mental health phenotypes

We used cross-trait LD-Score regression [46] to estimate the genetic correlation between lifetime cannabis use and 25 other traits using GWAS summary statistics. The genetic covariance is estimated using the slope from the regression of the product of z-scores from 2 GWASs on the LD score. The estimate represents the genetic covariation between the 2 traits based on all polygenic effects captured by SNPs. Summary statistics from well-powered GWASs were available for 25 relevant substance use and mental health traits, including nicotine, alcohol and caffeine use, schizophrenia, depression, bipolar disorder, and loneliness (Supplementary Table S5). To correct for multiple testing we adopted a Bonferroni corrected p-value threshold of significance of 0.002. LD scores were based on the HapMap 3 reference panel, restricted to European populations [47].

### Causal association between cannabis use and schizophrenia: Two-sample Mendelian randomisation

We performed two-sample Mendelian randomisation analyses (MR) [29] to examine whether there is evidence for a causal relationship from cannabis use to schizophrenia or vice versa. All analyses were performed with the R package of database and analytical platform *MR-Base* [*48*].

MR utilizes genetic variants strongly associated with an exposure variable as an ‘instrument’ to test for causal effects of the exposure on an outcome variable. This approach minimizes the risk of spurious findings due to confounding or reverse causation present in observational studies, provided that the following assumptions are met: 1) the genetic instrument is predictive of the exposure variable, 2) the genetic instrument is independent of confounders, and 3) the genetic instrument is not directly associated with the outcome variable, other than by its potential causal effect through the exposure (i.e. there is no directional pleiotropy) [49]. Two-sample Mendelian randomisation refers to the application of Mendelian randomisation methods to well-powered summary association results estimated in non-overlapping sets of individuals [29] in order to reduce instrument bias towards the exposure-outcome estimate.

Bi-directional causal effects were tested between lifetime cannabis use and schizophrenia. We used genetic variants from the cannabis GWAS as well as those from the largest schizophrenia GWAS [50] to serve as instruments (*gene-exposure association*). For lifetime cannabis use we used 2 genetic instruments; 1) an instrument including all independent genetic variants that were genome-wide significantly associated with lifetime cannabis use (p<5e-08; 4 SNPs), and 2) an instrument including independent variants with a more lenient significance threshold (p<1e-05; 37 SNPs). For schizophrenia we used one genetic instrument, including independent genetic variants that were genome-wide significantly associated with schizophrenia (instrument *p*<5e-08; 109 SNPs). Information on the included SNPs in the genetic instruments is provided in Supplementary Table S6.

Genetic variants were pruned (r^2^<0.01) and the remaining genetic variants (or proxies (r^2^≥0.8) when an instrumental SNP was not available in the other GWAS) were then identified in GWAS summary-level data of the outcome variable (*gene-outcome association*). Note that because not all exposure SNPs or their proxies are necessarily available also in the outcome dataset and because some SNPs were palindromic not all independent SNPs identified in the exposure dataset have been included in the analyses (see Supplementary Table S6).

Evidence for both a gene-exposure and a gene-outcome association suggests a causal effect, provided that the MR assumptions are met. To combine estimates from individual genetic variants we applied inverse-variance weighted (IVW) linear regression [51]. In addition, 4 sensitivity analyses more robust to horizontal pleiotropy were applied, each relying on distinct assumptions regarding instrument validity: Weighted Median [52], MR-Egger [30], Simple Mode, and Weighted Mode [53]. The weighted median approach provides a consistent estimate of the causal effect even when up to 50% of the weight comes from invalid instruments [52]. MR-Egger regression applies Egger’s test to MR instruments that consist of multiple genetic variants [30, 54]. MR-Egger provides a consistent estimate of the causal effect, provided that the strength of the genetic instrument (the association between SNPs and exposure) does not correlate with the effect the instrument has on the outcome (i.e. the InSIDE assumption (Instrument Strength Independent of Direct Effect)). This is a weaker assumption than the assumption of no pleiotropy. Finally, the Simple and Weighted Mode methods can produce an unbiased result, as long as the most common causal effect estimate is a consistent estimate of the true causal effect [the Zero Modal Pleiotropy Assumption (ZEMPA), 53].

To calculate variance explained (R^2^) by the instrument, first we selected a single SNP to obtain an estimate of the phenotypic variance, var(y). Assuming effect sizes are normally distributed, we used the quantile function of the student t-distribution to transform the p-value of the SNP association into an estimate of t, t^. The number of degrees of freedom and N were based on the effective sample size (4/(1/cases+1/controls)). The effective sample sizes were estimated at N=130,072 for schizophrenia and N=180,934 for cannabis use. The corresponding value of r was calculated using the formula t^=r / (sqrt[(1− R^2^)/(N−2)] and obtained the R^2^ that corresponds to t^ with the online tool http://vassarstats.net/rsig.html. Subsequently, we approximated the variance of the phenotype y using var (y)=(2*MAF*(1-MAF)*β^2^)/ R^2^ in which MAF denotes the Minor Allele Frequency and β the effect size of the specific SNP. Finally, we used the estimated value of var (y) to calculate the R^2^ for the remaining SNPs of interest using R^2^=(2*MAF*(1−MAF)*β^2^)/var(y); and summed the R^2^ of all SNPs of interest included in the instrumental variable to obtain an estimate of the total R^2^ explained by the instrument.

## Acknowledgements

JP and JMV are supported by the European Research Council [Beyond the Genetics of Addiction ERC-284167, PI JM Vink]. NAG is supported by US National Institutes of Health, National Institute on Drug Abuse R00DA023549. JLT is supported by the Netherlands Organization for Scientific Research (NWO; Rubicon grant 446-16-009). SM is supported by an Australian Research Council Fellowship. Statistical analyses were partly carried out on the Genetic Cluster Computer (*http://www.geneticcluster.org*) hosted by *SURFsara* and financially supported by the *Netherlands Organization for Scientific Research* (NWO 480-05-003 PI: Posthuma) along with a supplement from the Dutch Brain Foundation and the VU University Amsterdam. MGN is supported by Royal Netherlands Academy of Science Professor Award to DIB (PAH/6635). Part of the computation of this project was funded by NWO exact sciences for the application: “Population scale Genetic Analysis” awarded to MGN. The genome-wide association analysis on the UK-Biobank dataset has been conducted using the UK Biobank resource under application numbers 9905, 16406 and 25331.

We would like to thank the research participants and employees of 23andMe for making this work possible. JM’s contributions were partially supported by the Peter Boris Chair in Addictions Research. SS-R was supported by the Frontiers of Innovation Scholars Program (FISP; #3-P3029), the Interdisciplinary Research Fellowship in NeuroAIDS (IRFN; MH081482) and a pilot award from DA037844. The Estonian Genome Center, University of Tartu was supported by the European Union through the European Regional Development Fund (Project No. 2014-2020.4.01.15-0012) and the European Union’s Horizon 2020 research and innovaLon programme under grant agreements No 692065 and 692145. JK was supported by Academy Professorship grants by the Academy of Finland (263278, 292782). MR is a recipient of a Miguel de Servet contract from the Instituto de Salud Carlos III, Spain (CP09/00119 and CPII15/00023). Her work was co-financed Instituto de Salud Carlos III (PI16/01505, PI17/00289), and by the European Regional Development Fund (ERDF), the Agència de Gestió d’Ajuts Universitaris i de Recerca-AGAUR, Generalitat de Catalunya, Spain (2014SGR1357), the NARSAD Young Investigator Grant from the Brain & Behavior Research Foundation and the European Union H2020 Programme (H2020/2014-2020) under grant agreements Nos. 667302 (CoCA).

## Author contributions

KJHV, EMD, and JMV were responsible for the study concept and the design of the study. ICC contributed existing genome-wide summary data from the International Cannabis Consortium. SS, SS-R, MGN, BMLB, J-SO, HI, MDZ, MB, FRD, PF, SM, JRBP, AAP, and DP performed or supervised genome-wide association analyses. JAP performed the quality control and meta-analysis of genome-wide association studies, under supervision of KJHV, BMLB, and JMV. JAP, KJHV, ZG, JLT, AA, MRM, and EMD contributed to secondary analyses of the data. JAP, KJHV, ZG, NAG, EMD, and JMV wrote the manuscript. JLD, SJTB, CAH, ACH, PACL, PAFM, RM, WM, GWM, AJO, ZP, JARQ, TP, MR, JK, MPMB, JTB, TDS, JG, DIB, and NGM contributed to data acquisition of the samples in the International Cannabis Consortium. SLE, HDW, LKD, and JMK contributed to data acquisition and analysis for the 23andMe dataset. All authors provided critical revision of the manuscript for important intellectual content.

### Contributors to the 23andMe Research Team

M. Agee, B. Alipanahi, A. Auton, R.K. Bell, K. Bryc, S.L. Elson, P. Fontanillas, N.A. Furlotte, D.A. Hinds, K.E. Huber, A. Kleinman, N.K. Litterman, J.C. McCreight, M.H. McIntyre, J.L. Mountain, E.S. Noblin, C.A.M. Northover, S.J. Pitts, J. Fah Sathirapongsasuti, O.V. Sazonova, J.F. Shelton, S. Shringarpure, C. Tian, J.Y. Tung, V. Vacic, and C.H. Wilson

### Contributors to the International Cannabis Consortium

S. Stringer, C.C. Minică, K.J.H. Verweij, H. Mbarek, M. Bernard, J. Derringer, K.R. van Eijk, J.D. Isen, A. Loukola, D.F. Maciejewski, E. Mihailov, P.J. van der Most, C. Sánchez-Mora, L. Roos, R. Sherva, R. Walters, J.J. Ware, A. Abdellaoui, T.B. Bigdeli, S.J.T. Branje, S.A. Brown, M. Bruinenberg, M. Casas, T. Esko, I. Garcia-Martinez, S.D. Gordon, J.M. Harris, C.A. Hartman, A.K. Henders, A.C. Heath, I.B. Hickie, M. Hickman, C.J. Hopfer, J.J. Hottenga, A.C. Huizink, D.E. Irons, R.S. Kahn, T. Korhonen, H.R. Kranzler, K. Krauter, P.A.C. van Lier, G.H. Lubke, P.A.F. Madden, R. Mägi, M.K. McGue, S.E. Medland, W.H.J. Meeus, M.B. Miller, G.W. Montgomery, M.G. Nivard, I.M. Nolte, A.J. Oldehinkel, Z. Pausova, B. Qaiser, L. Quaye, J.A. Ramos-Quiroga, V. Richarte, R.J. Rose, J. Shin, M.C. Stallings, A.I. Stiby, T.L. Wall, M.J. Wright, H.M. Koot, T. Paus, J.K. Hewitt, M. Ribasés, J. Kaprio, M.P.M. Boks, H. Snieder, T. Spector, M.R. Munafò, A. Metspalu, J. Gelernter, D.I. Boomsma, W.G. Iacono, N.G. Martin, N.A. Gillespie, E.M. Derks, and J.M. Vink

## Conflict of interest

PF, SLE, and members of the 23andMe Research Team are employees of 23andMe Inc. JARQ was on the speakers’ bureau and/or acted as consultant for Eli-Lilly, Janssen-Cilag, Novartis, Shire, Lundbeck, Almirall, BRAINGAZE, Sincrolab, and Rubió in the last 5 years. He also received travel awards (air tickets + hotel) for taking part in psychiatric meetings from Janssen-Cilag, Rubió, Shire, and Eli-Lilly. The Department of Psychiatry chaired by him received unrestricted educational and research support from the following pharmaceutical companies in the last 5 years: Eli-Lilly, Lundbeck, Janssen-Cilag, Actelion, Shire, Ferrer, and Rubió.

